# Dynamics of Allosteric Transitions in Dynein

**DOI:** 10.1101/306589

**Authors:** Yonathan Goldtzvik, Mauro L. Mugnai, D. Thirumalai

**Affiliations:** Department of Chemistry, University of Texas at Austin, Austin, TX 78712

## Abstract

Cytoplasmic Dynein, a motor with an unusual architecture made up of a motor domain belonging to the AAA+ family, walks on microtubule towards the minus end. Prompted by the availability of structures in different nucleotide states, we performed simulations based on a new coarse-grained model to illustrate the molecular details of the dynamics of allosteric transitions in the motor. The simulations show that binding of ATP results in the closure of the cleft between the AAA1 and AAA2, which in turn triggers conformational changes in the rest of the motor domain, thus poising dynein in the pre-power stroke state. Interactions with the microtubule, which are modeled implicitly, substantially enhances the rate of ADP release, and formation of the post-power stroke state. The dynamics associated with the key mechanical element, the linker (LN) domain, which changes from a straight to a bent state and vice versa, are highly heterogeneous suggestive of multiple routes in the pre power stroke to post power stroke transition. We show that persistent interactions between the LN and the insert loops in the AAA2 domain prevent the formation of pre-power stroke state when ATP is bound to AAA3, thus locking dynein in a non-functional repressed state. Motility in such a state may be rescued by applying mechanical force to the LN domain. Taken together, these results show how the intricate signaling dynamics within the motor domain facilitate the stepping of dynein.

## 2 Introduction

Molecular motors are spectacular nanomachines that transport cargos by moving processively along filamentous actin and microtubules (MT).^1–3^ Of the three motor families, kinesins and dyneins walk along MTs whereas myosins step on filamentous actin. Although a number of questions still remain unanswered, it is fair to say that the mechanism of processive motion of kinesin-1 and myosin V is fairly well understood, thanks to remarkable experimental studies conducted over the last twenty five years.^2, 4–16^ In contrast, the characterization of the motility of cytoplasmic dynein has lagged behind. Although discovered over fifty years ago,^17^ it is only in the last decade the complexity and variations in the architecture of dyneins have been revealed, raising various questions that challenge our views of information processing in nanoscale biological machines. Even though this issue, which relates to to allosteric communication in protein complexes, is most interesting, our focus here is to provide quantitative insights into the molecular basis of the dynamics of allosteric transitions between distinct functional states in the motility of dynein, a most intriguing molecular motor.

Cytoplasmic dynein transports cargo along MTs, and is a minus end-directed motor. Among dyne’s different cargoes are endosomes, lysosomes, and mitochondria.^18–20^ Additionally, cytoplasmic dynein (or dynein from now on) is involved in the process of positioning of the spindle during cell division.^21^ In almost all respects, dynein is distinct in comparison with other cytoskeletal motors. (i) In contrast to conventional kinesin, which transports cargo towards the + end of the polarized MT, dynein steps towards the - end of the polar track. (ii) The motor domain belongs to the class of AAA+ family.^22^ Therefore, it stands to reason that dynein motors must have evolved from a different lineage compared to myosin and kinesin. (iii) Several of the well studied AAA+ enzymes, such as bacterial chaperonin GroEL and protein degradation machines, are oligomeric assemblies often consisting of identical domains. The motor domain in dynein, on the other hand, is a hetero-oligomer with six AAA domains, originating from a single polypeptide chain. (iv) Unlike many other AAA+ assemblies, nucleotide activity in each nucleotide binding cite is distinct. How the diverse changes in the nucleotide chemistry affect dynein motility is still largely unknown. (v) The size of the dynein motor domain (MD), is significantly larger than the MD of kinesin and myosin. The length of the MD of dynein along its longest axis is about 25 nm, in contrast to the diameter of kinesin, which is only about 5-6 nm.^23, 24^ (vi) The complex MD of dynein comprises of several appendages that are unique among cytoskeletal motors and are crucial to the proper functioning of dynein.^23, 25, 26^ Owing to the presence of these peculiar structural elements, the connection between the architecture of the MD and dynein motility is not transparent, as is the case for example in myosin V.^27^

The core of the MD consists of a hexameric AAA+ (AAA1-AAA6) assembly, covalently linked together (Fig. 1). Of these, the first four AAA domains of dynein can catalyze ATP to a varying degree of efficiency. However, it has been shown that the AAA1 domain is the main ATP catalytic site of the motor.^28^ Other than the AAA domains, the structure of dynein contains several prominent elements (Fig. 1), among which is the LN, a large and elongated domain, spanning the diameter of the entire AAA+ ring (shown in purple in Fig. 1). The LN is involved in the power stroke, the conformational transition in the motor that is responsible for propelling the motor forward to the minus end of the MT.^29, 30^ An interesting structural feature of dynein is that the microtubule binding domain (MTBD) is located at the end of the stalk, which is approximately 10 nm long coiled coil structure emerging from the AAA4 domain (shown in yellow in Fig. 1). The MTBD is spatially well-separated from the main ATP hydrolysis site in the AAA1 domain, which raises the question of how the two sites communicate. In order for the ATP binding site and the MTBD to communicate, an elaborate set of conformational changes has to occur.^23, 25, 31, 32^ While the details of this process are still being worked out, the most noticeable conformational change is a shift in the relative position of the coils in the stalk.^31, 33, 34^ This transitions appears to be induced by a structural element, referred to as the strut (orange element in Fig. 1).

**Figure 1:**
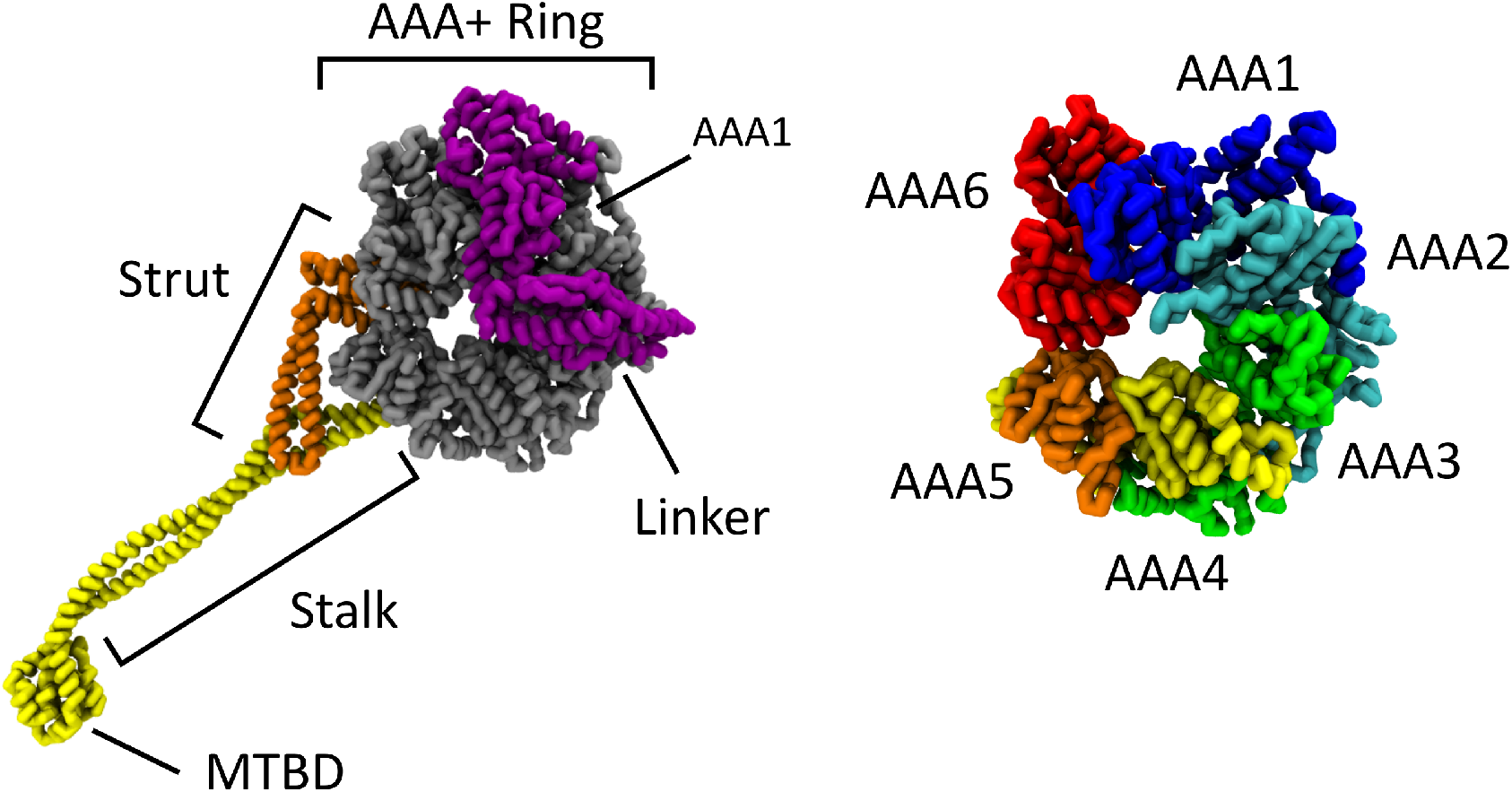
Key structural elements of the dynein motor domain (MD). Left: Parts list of the MD: hexameric AAA+ ring (**gray**), linker (**purple**), strut (**orange**), stalk and Microtubule Binding Domain (MTBD) (**yellow**). Right: The AAA+ ring, comprised of the six AAA domains, is assembled from a single polypeptide chain: AAA1 (**blue**), AAA2 (**cyan**), AAA3 (**green**), AAA4 (**yellow**), AAA5 (**orange**), and AAA6 (**red**).

Just like other motors, dynein motility depends on the ability of the MDs to translate the binding and hydrolysis of ATP into mechanical work. However, the ability to hydrolyze ATP by AAA1-AAA4 (AAA5 and AAA6 do not bind ATP) varies greatly, and their relative contributions to motility also differ considerably.^28, 35–37^ Binding of ATP to AAA1 (with ADP in AAA3) induces rapid dissociation of the motor from the MT, and a priming stroke of the LN, leading to a pre-power stroke state in which the LN is in a bent conformation (upper row in Fig. 2).^29, 36, 38, 39^ In addition, ADP release from AAA1 is accelerated when the motor binds to MT, just as in kinesin.^40^ Interestingly, the binding of the motor to MT also accelerates the power stroke transition in the LN, leading to a post-power stroke state (lower row of Fig. 2).^41^ This suggests that the process of ADP release and the power stroke must be coupled. What makes dynein so perplexing is that the allosteric sites, the ATP binding site in AAA1 and the MTBD, are located far from each other (Fig. 1). It is clear that in order for dynein to walk processively on MT, an elaborate scheme of allosteric control and conformational transitions in the MD has to take place, and the resulting signal has to be transmitted to the MTBD.

**Figure 2:**
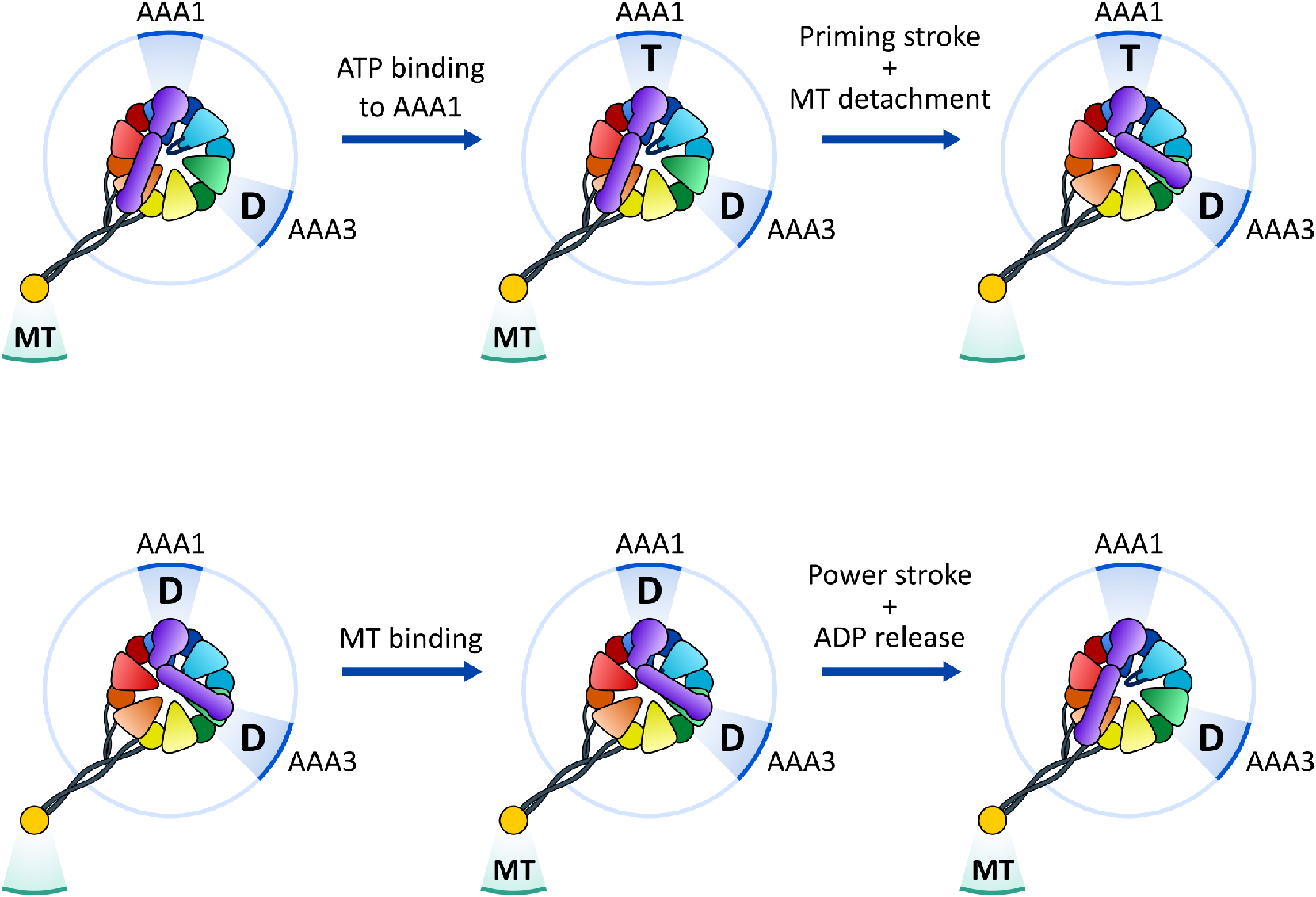
Mechanochemical reaction cycle pathways in dynein. Starting in the MT bound *apo* state, with ADP in the AAA3 site, ATP binds to the AAA1 site. ATP binding induces detachment of dynein from the MT as well as the priming stroke in the LN domain (**upper row**). Upon ATP hydrolysis and phosphate release the MD has ADP in AAA1. Dynein then binds to the MT, which in turn accelerates the released of ADP and the power stroke motion in the LN (**lower row**).

A structural picture of how these allosteric transitions occur is starting to emerge over the last few years, thanks to the availability of structures from different species in several allosteric states.^23, 25, 26, 36^ In order to go beyond the structures and provide the molecular basis of the dynamics of allosteric communication and gating in dynein, we performed Brownian dynamics simulations, using a new coarse-grained version of the Self-Organized Polymer (SOP) model^42^ (summarized in **Experimental Procedures** with technical details in the Supplementary information (SI)) to monitor the dynamical changes that occur in the MD in response to either ATP or MT binding.^43, 44^ In general, coarse-grained models have been remarkable insights into the dynamics of motors.^45–47^ Using these simulations, we consider the following questions inspired by several experimental findings. What is the molecular mechanism of long-range structural transitions underlying the ATP/MT binding induced priming and power stroke? How does the AAA3 unit modulate the allosteric communication in the motor? What could be the molecular basis of gating in dynein? The results of our simulations provide answers to these questions and shed light on the intricate allostery in dynein. Our simulations reproduce a range of known aspects of dynein allostery, such as the structural changes in the MD that occur in response to the binding of ATP and MT. In particular, the simulations show that, dynamically, the formation of the priming stroke (LN in the bent state) occurs first upon closing of the cleft between AAA1 and AAA2 followed by changes in AAA5, AAA6, and the stalk. Furthermore, our data suggests that MT binding accelerates the execution of the power stroke and ADP release. We show that the AAA2 domain is involved in the suppression of dynein activity when ATP is bound to AAA3. Our simulations show that the persistent interactions of the insert loops (ILs) with the LN domain when ATP is bound to AAA3 results in the repression of dynein motility. Application of mechanical force to the LN domain restores motility, as demonstrated in single molecule experiments,^37^ of the repressed state by facilitating the allosteric changes that occur in the normal catalytic cycle. Based on this work we make testable predictions, especially in regard to the importance of interactions between IL and the LN domain in dynein motility.

## 3 Results

In order to answer the questions posed above, we took a computational approach using the following strategy: (i) Construct a tractable model of the dynein MD, with which we can account for responses to external inputs such as the binding of ATP or interactions with MT. (ii) Require that the simulations using the model reproduce the experimentally observed response of the motor to ligand or ATP binding. (iii) Introduce and elucidate the repressing effects of ATP binding to the AAA3 domain. (iv) Analyze the consequences of the external force, applied to the MD.

### Allosteric Communication Between the ATP Binding Site and the MT

Several studies have shown that binding of ATP to the primary ATP binding site between the AAA1 and AAA2 domains (referred to as AAA1/2) induces a pronounced conformational transition from a straight to bent state in the LN domain (priming stroke), leading to detachment of the motor from the MT.^29, 30, 41^ Furthermore, once ATP hydrolysis and phosphate (Pi) release are complete, binding of dynein to MT accelerates the release of ADP, resulting in the conformational transition from the bent to the straight state in the LN (power stroke).^38^ We require that our model reproduce the global responses of dynein to ATP and MT binding.

It is suspected that allosteric communication between the AAA1/2 ATP binding site and the MTBD is transmitted through a sliding motion in the coiled coil stalk domain (shown in yellow in Fig. 1).^31, 33, 48^ How the information is transferred between AAA1/2 and the stalk, which are spatially well separated, is less clear. From the architecture of the motor, it is reasonable to argue that the allosteric communication must occur via conformational changes in the AAA5 and AAA6 domains. To support this argument, we rely on two pieces of evidence. First, structural studies of dynein suggest that the strut, a domain that connects the AAA5 and stalk domains (orange element on the left in Fig. 1), is involved in the conformational changes in the stalk.^34^ Second, when comparing crystal structures of dynein (taken from different species) in the ATP bound and no nucleotide (*apo*) states, setting aside the LN and stalk domains, most of the conformational changes occur in the AAA1, AAA5, and AAA6 domains (see Fig. S1).^23, 25, 26^ Therefore, we assume, when constructing our model, that the allosteric communication pathway between AAA1/2 and the MTBD involves the participation of the AAA5, AAA6, and the stalk. We group these domains together and refer to them as AAA5/6/S.

In order to ensure that our results are consistent with experimental observations, which would serve as a validation of the model and the simulation strategies, we considered two scenarios that naturally occur during the catalytic cycle of dynein. In scenario I, starting from the *apo* state, ATP binds to the AAA1 domain, inducing a cascade of conformational changes along the AAA5/6/S allosteric pathway, resulting in the detachment of the MD from the MT. In this process, the LN undergoes a priming stroke (Fig. 2) from a straight (S) to bent (B) conformation (*LN_S_* → *LN_B_* transition). In scenario II, we assume that the motor has ADP in the AAA1 domain, thus creating the pre-power stroke state. Dynein then binds to MT at the MTBD, accelerating the release of ADP, resulting in a power stroke (*LN_B_* → *LN_S_* transition) in the LN domain. These transitions are schematically illustrated in Fig. 2.

#### Conformational Transitions upon ATP Binding

In order to probe the dynamical transitions that occur upon binding of ATP in scenario I, we probed the time-dependent conformational changes that occur in the AAA1/2 ATP binding domain during the transition from the *apo* state (referred to as state E) to the ATP bound state (state A). The most notable conformational change at the AAA1/2 site is the closure of the cleft between the AAA1 and AAA2 domains (Fig. 3). The decrease in Δ_*N*1714–*S*2065_, the distance between residue N1714 in AAA1 and residue S2065 in AAA2 (upper left panel in Fig. 4a and Fig. 4b), with time shows the closure of the cleft between these two domains upon ATP binding. In contrast, the cleft remains open in the *apo* state (upper right panel in Fig. 4a).

**Figure 3:**
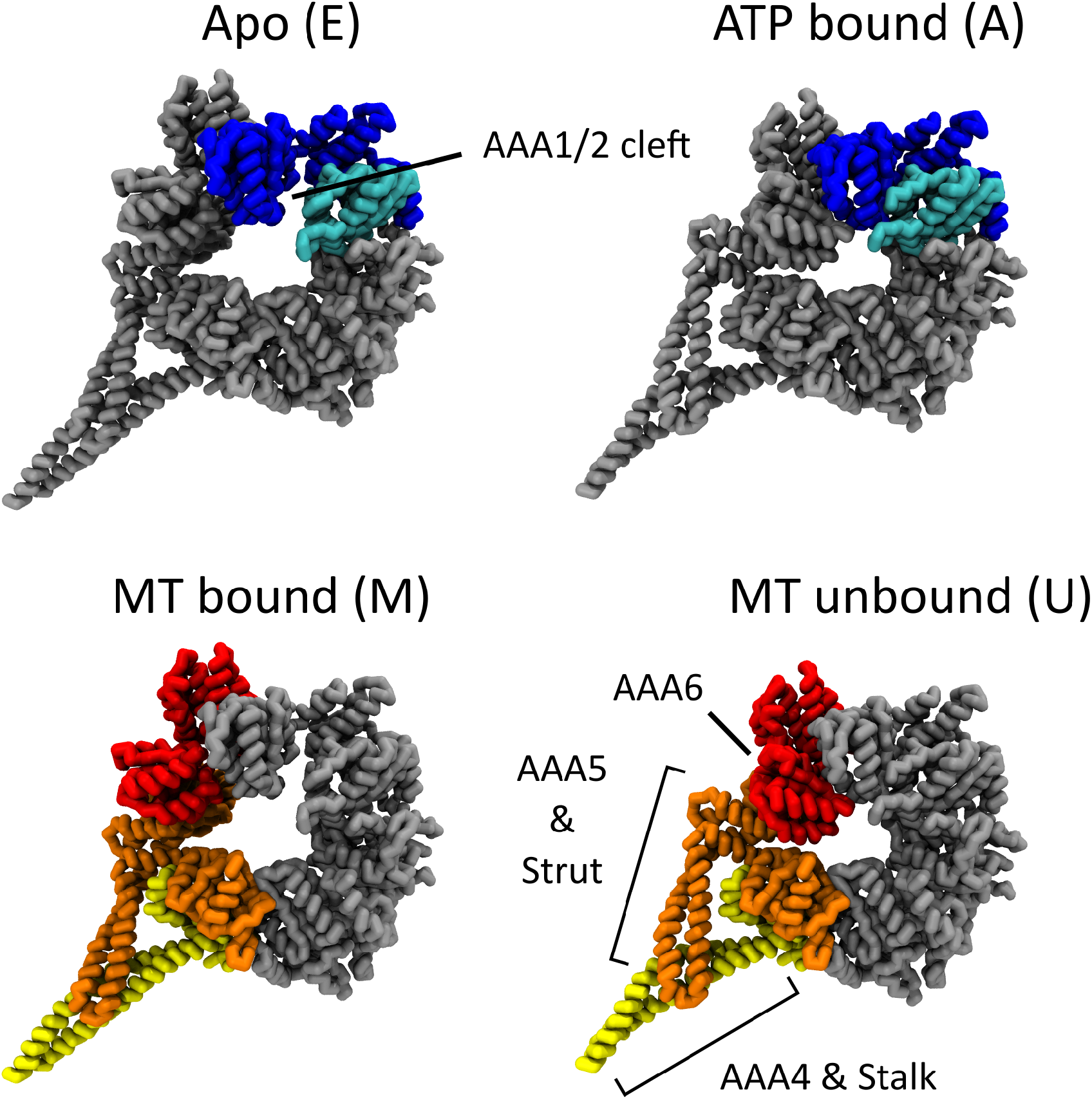
Conformational transitions in dynein. Top: Structures of dynein in the *apo* (E) state (left) or ATP bound (A) state (right). The AAA1 and AAA2 units are marked in blue and cyan respectively. The cleft between the AAA1 and AAA2 domains is open in the E state but is closed in the A state. Bottom: Structures of dynein in the MT bound (M) state (left) and MT unbound (U) state (right). The AAA5, AAA6, and stalk domains are highlighted in orange, red, and yellow respectively. Notice the conformational differences between the two states, particularly in the strut sub-domain.

**Figure 4:**
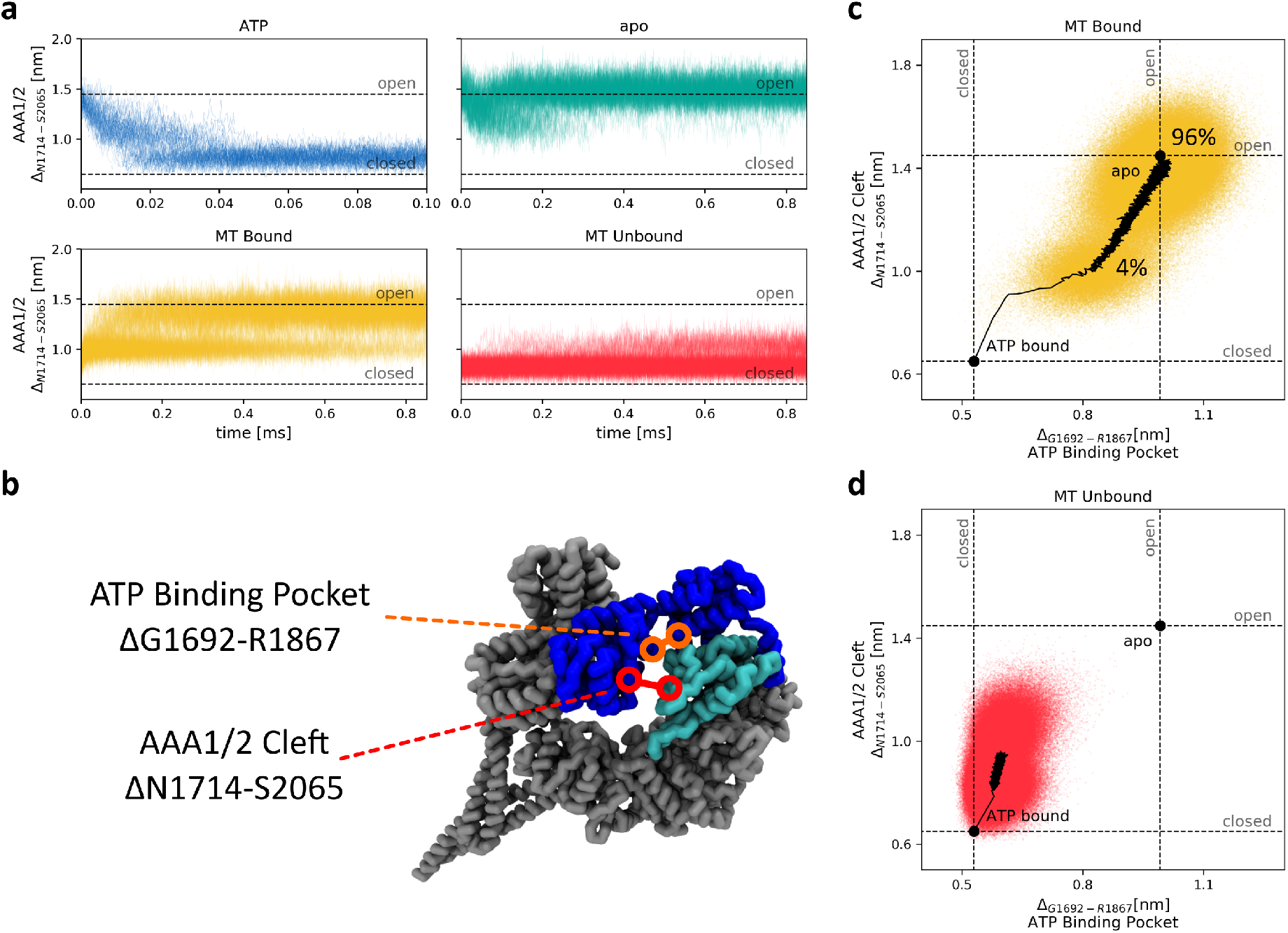
Consequences of dynein interactions with ATP and MT. (**a**) The value Δ_*N*1714–*S*2065_ (AAA1/2 cleft) is plotted as a function of time in 100 trajectories. The dashed lines mark the values of Δ_*N*1714–*S*2065_ in the A (closed) and E (open) states. (**blue**) The AAA1/2 cleft closes rapidly in response to the binding of ATP. (**green**) The cleft remains open in the absence of nucleotide. (**yellow**) MT binding accelerates the opening of the AAA1/2 cleft and ADP release. (**red**) In the absence of MT, opening of the cleft is slow. (**b**) The extent of conformational change in the AAA1/2 ATP binding pocket is monitored by Δ_*G*1692–*R*1867_ (**orange**) while Δ_*N*1714–*S*2065_ (**red**) measures whether the AAA1/2 cleft is open or closed. (**c**) Plot of Δ_*N*1714–*S*2065_ (AAA1/2 cleft) against Δ_*G*1692–*R*1867_ (ATP binding pocket). The distances are sampled in 100 trajectories when dynein is bound to MT (**yellow**). Both the AAA1/2 cleft and the ATP binding pocket are either partially or completely open. The percentage of fully/partially open clefts/ATP binding pockets, sampled during the second half of the simulation, is shown by the numbers in the plot. The mean trajectory, averaged over all trajectories, is shown by the black curve. (**d**) Same as (**c**) except the distances Δ_*N*1714–*S*2065_ (AAA1/2 cleft) versus Δ_*G*1692–*R*1867_ (ATP binding pocket) are shown in the absence of MT (**red**). While the AAA1/2 cleft partially opens, the ATP binding pocket remains predominantly closed. The dashed are the values of Δ_*N*1714–*S*2065_ and Δ_*G*1692–*R*1867_ in the A (closed) and E (open) states.

Binding of ATP to the AAA1 domain leads to a cascade of conformational changes across the entire motor domain.^23, 25, 26, 36, 49^ We probed the dynamics associated with transitions in the AAA5/6/S domains due to nucleotide binding to AAA1. In order to account for the decreased affinity for MT, due to ATP binding to AAA1, we monitored the transition of the AAA5/6/S domains from the MT bound state (referred to as state M) to the unbound state (referred to as state U). However, the transition between the M and U states was allowed in the simulations only after the distance Δ_*N*1714–*S*2065_ decreased below 0.8 nm. This procedure mimics the inference from experiments that the allosteric transitions occur in AAA5/6/S only as a consequence of the conformational changes at the AAA1/2 site.^23, 25, 36, 38^ We tracked the allosteric transition by monitoring Δ_*N*1767–*N*3709_, the distance between residues N1767 and N3709 (Figs. S2). Comparison of the results in the top and bottom left panels in Fig. S2a shows that the AAA5/6/S domains transition from the conformations in the M state to the one adopted in the U state occurs only after the closure of the cleft. In contrast, the right panels in Fig. S2a show that this transition does not take place when the cleft remains open, thus confirming that the M to U transition is a direct result of the cleft closure.

In order to ensure that the transitions described above are a direct result of ATP binding and not an artifact of the simulation methodology, we performed additional simulations of dynein in the *apo* state. In these simulations the AAA1/2 was restrained to be in the E state with the cleft open (Fig. 3). We allowed for the transition in the AAA5/6/S domains if Δ_*N*1714–*S*2065_ is less than 0.8 nm. Fig. S2 shows that in the simulations of dynein with ATP, the AAA1/2 site changed conformations, which in turn affects the conformations in AAA5/6/S. On the other hand, in the simulations of the *apo* state, no such transition occurred (see the upper right panel in Fig. S2a). Thus, the proposed CG model agrees with structural and kinetic experiments, which pointed out that the conformational changes in AAA5/6/S take place only upon ATP binding to AAA1/2.^23, 25, 38^

#### Transitions resulting from MT binding

In scenario II, dynein binds to MT at the MTBD. In our simulations we do not include the MTBD explicitly but we assume that when bound to MT, the AAA5/6/S domains are predominantly in the M state. By stabilizing the M state with respect to the U state at the AAA5/6/S domains, we implicitly include the effects of MT binding (see SI for details of the simulation procedure). The AAA1/2 is in the ATP bound A state at the start of the simulations but the stabilizing interactions at this site are weakened (see SI) in order to reflect that the binding site contains ADP and not ATP. If Δ_*N*1714–*S*2065_, reporting the fate of the cleft, increases beyond 1.1 nm, the transition to the nucleotide free E state is facilitated.

Comparison of the M and U states show that binding to MT significantly accelerates the opening of the cleft at the AAA1/2 site (compare the two lower panels in Fig. 4a), which is consistent with the observation made in experiments.^41^ Furthermore, while there are fluctuations in the AAA1/2 cleft in the simulations without the effects of MT, the cleft does not open fully within the simulation time. Although Δ_*N*1714–*S*2065_ is a good measure of the opening of the AAA1/2 cleft, it is not necessarily an appropriate indicator of what transpires at the ATP binding site itself. In order to monitor the conformational changes at the ATP binding site directly, we measured Δ_*G*1692–*R*1867_, the distance between residues G1692 and R1867, located at the ATP binding site (Fig. 4b). We tracked Δ_*N*1714–*S*2065_ and Δ_*G*1692–*R*1867_ simultaneously, both in the M and U states. Fig. 4c shows that when the AAA5/6/S domains adopted the conformations of the MT bound M state, both the AAA1/2 cleft and ATP binding pocket are open. In contrast, in the absence of MT, while there is partial opening of the AAA1/2 cleft, the ATP binding pocket itself remains closed (Fig. 4d). This finding supports the hypothesis that conformational changes involving AAA5/6/S, facilitated by binding to the MT, are responsible for the acceleration of ADP release from the AAA1/2 site.^32, 50, 51^

### Conformational Transitions in the LN Domain

In addition to the regulation of affinity of dynein for MT, ATP binding and hydrolysis also control the conformations of the LN domain. In particular, it has been suggested that binding of ATP at the AAA1/2 site leads to a priming stroke in the LN domain.^52^ On the other hand, binding to MT accelerates the power stroke motion in the LN domain.^41^ To test whether our simulations support these findings, and to provide a structural view of the dynamics associated with these processes, we tracked the motion of the LN domain in both scenarios I and II.

#### Scenario I

In order to monitor the conformational changes in the LN domain to the binding of ATP, we measured Δ_*S*1315–*D*3297_, the distance between residue S1315 near the N-terminus of the LN domain and residue D3297 in the AAA5 domain (Fig. 5). Both the residues are at the interface between the LN and the AAA5 domains (see the static structures in Fig. 5), making Δ_*S*1315–*D*3297_ a reasonable order parameter for assessing if the LN domain is in the post-power stroke conformation or whether the LN domain is detached from the AAA5 binding site.

**Figure 5:**
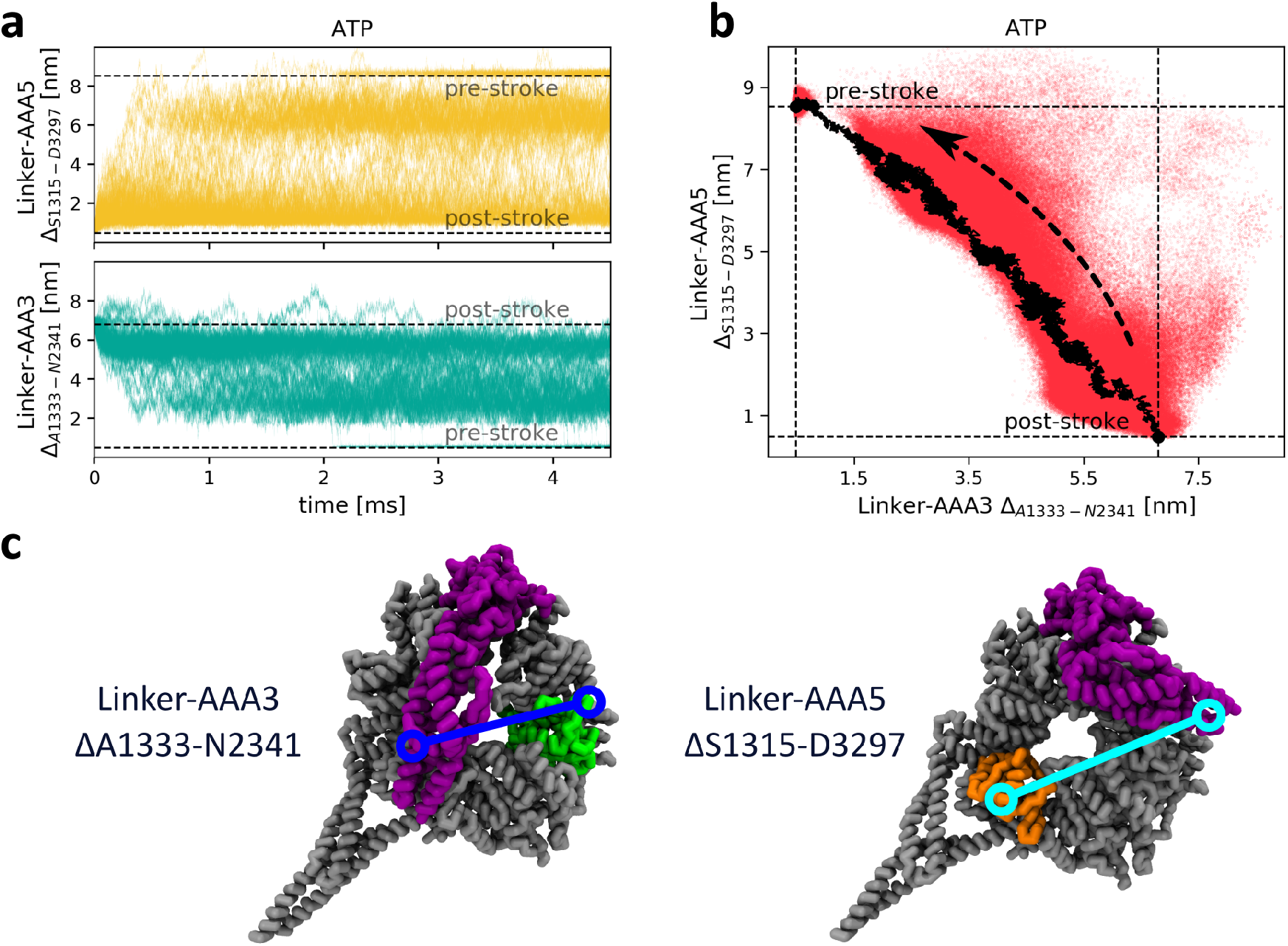
The priming stroke of dynein. (**a**) Response of the LN to ATP binding as a function of time is tracked using Δ_*S*1315–*D*3297_ (LN-AAA5 distance) and Δ_*A*1333–*N*2341_ (LN-AAA3 distance), with the dashed lines being the values of Δ_*S*1315–*D*3297_ and Δ_*A*1333–*N*2341_ in the pre- and post-power stroke states. When dynein binds ATP, the LN detaches from the AAA5 domain (**yellow**). The distance between the LN and the AAA3 decreases in some of the trajectories until the LN reaches the pre-power stroke conformation (**green**). There is a great deal of heterogeneity, which is reflected in the large variations from one trajectory to another. (**b**) Scatter plot of Δ_*S*1315–*D*3297_ versus Δ_*A*1333–*N*2341_ in the presence of ATP. The LN diffuses between many partially bent conformations before completing the priming stroke (**red**). The black curve shows the average of 13 trajectories in which the LN reaches the AAA3 binding site. The arrow indicates the general direction of the priming stroke transition. (**c**) The distances between the LN domain and the AAA3 and AAA5 domains are monitored using Δ_*A*1333–*N*2341_ (**blue**) and Δ_*S*1315–*D*3297_ (**cyan**), respectively. The mean trajectory, averaged over the 13 trajectories in which the LN reaches the AAA3 binding site, is shown by the dark curve.

The upper panel in Fig. 5a shows that in a large percentage of the trajectories the LN domain detaches from the AAA5 domain, as can be seen from an increase in Δ_*S*1315–*D*3297_. To ensure that this is a result of ATP binding, we also measured Δ_*S*1315–*D*3297_ in simulations of the *apo* state with no ATP bound at the AAA1/2 site. With the exception of a couple of trajectories, the LN domain remained bound to the AAA5 domain (Fig. S3). In order to ascertain if the LN domain reaches the AAA3 binding site in the pre-power stroke state, we measured Δ_*A*1333–*N*2341_, the distance between residue A1333 in the LN domain and residue N2341 in the AAA3 domain. These residues come into contact only in the pre-power stroke, *LN_B_* state, and therefore Δ_*A*1333–*N*2341_ is an indicator of whether the priming stroke is complete. The lower panel in Fig. 5a shows that in a number of trajectories, due to ATP binding to AAA1, the LN domain reaches *LN_B_* state within the simulation time. It should be noted that this transition is not complete in most trajectories, at least within the time scale of our simulations. It is again worth remarking that there is a great deal of variation from trajectory to trajectory, underscoring the importance of heterogeneity associated with the transition involving the LN domain.

To better characterize the correlated motion in the dynamics of the priming stroke transition, we plotted Δ_*S*1315–*D*3297_ versus Δ_*A*1333–*N*2341_, obtained from a number of trajectories (Fig. 5b). As can be seen from both Figs. 5a and 5b, the LN detached from AAA5 rapidly. However, the binding to AAA3, indicated by the approach of Δ_*A*1333–*N*2341_, to its equilibrium value in the pre-power stroke, lags behind. Instead, the LN makes multiple transitions between the *LN_S_* and *LN_B_* states. In order to monitor the conformations of the LN itself we calculated *θ*_*LN*_, the angle between residues Q1255, F1418, and G1585, which directly measures the bending of the LN (Fig. S4). Interestingly, the LN does not bind to AAA3 and complete the priming stroke until relatively late in the transition. As can be seen in Fig. S4a (top panel), the LN spends a significant amount of time in a partially bent state. During the second half of the simulation (2.25 – 4.5*ms*) the LN has a probability of 44% to be only partially bent and 10% to be fully bent (Fig. S4a).

The picture of how the post-power stroke (*LN_S_*) state is destabilized by ATP binding is complex. Our simulations suggest that the LN dynamically comes into contact with the AAA2 insert loops (IL) in the post-power stroke state (Fig. 6). The interaction between the LN and the ILs in AAA2 leads to partial detachment of the LN from the AAA5 binding site even in the *apo* state (Fig. 6a). This is not entirely surprising, because in the crystal structure by Kon *et al*, the LN domain interacts with the IL in the ADP bound state (which is structurally similar to the *apo* state).^26^ The LN is also somewhat shifted from its AAA5 bound position in this structure, and is located in the cleft between AAA4 and AAA5 domains. Naively, this may suggest that favorable interactions between the LN and the AAA2 domains are not compatible with LN-AAA5 docking. Nevertheless, these interactions seem to stabilize the post stroke conformation as a whole (Fig. 6a and 6b). To further support this claim, we performed mutation simulations of the *apo* state in which the stabilizing interactions between the LN and the AAA2 domain are switched off. Fig. S5 shows that without the stabilizing interactions between the LN and the ILs, the post-power stroke conformation is destabilized as the LN is more likely to detach from the AAA5 domain, as indicated by the large values of Δ_*S*1315–*D*3297_, the distance reporting the LN-AAA5 interactions. In order to make a quantitative comparison between the wild type and mutation simulations in the *apo* state, we calculated the percentage of trajectories in which the LN was completely detached from both AAA2 and AAA5. Our calculations show that during the second half of the the mutation simulation, the LN has ~ 41% chance of being completely detached, in contrast to the wild type simulations in which the probability was only ~ 1%. This confirms our conclusion that the IL interactions with the LN stabilize the post-power stroke state.

**Figure 6:**
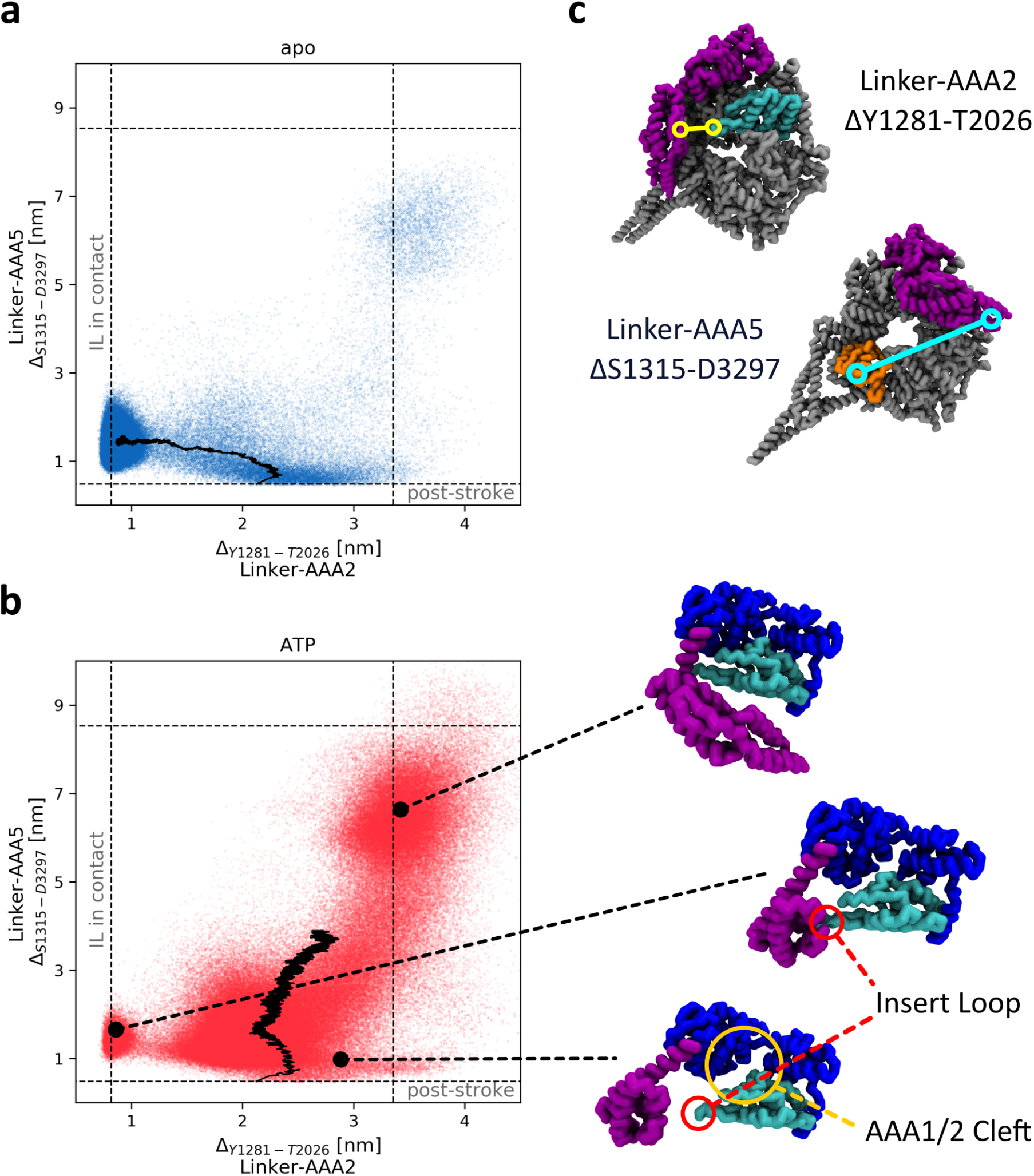
The effects of the Insert Loops (ILs) of AAA2 on the stability of the post-power stroke state. (**a**) Changes in Δ_*S*1315–*D*3297_ (LN-AAA5 distance) against Δ_*Y*1281–*T*2026_ (LN-AAA2 distance) sampled in 100 trajectories of dynein in the *apo* state (**blue**). In the absence of ATP, the LN still binds to the AAA2 ILs, suggesting that these interactions stabilize the post stroke conformations. The Δ_*S*1315–*D*3297_ and Δ_*Y*1281–*T*2026_ values in the reference structures are given by the dashed lines. The mean trajectory, averaged over all trajectories, is shown by the black curve. (**b**) Same as (**a**) except Δ_*S*1315–*D*3297_ (LN-AAA5 distance) against Δ_*Y*1281–*T*2026_ (LN-AAA2 distance) is shown in the ATP bound state (**red**). Before the LN detaches from the AAA5 domain, the the ILs in AAA2 tend to form contacts with the LN. Eventually the LN detaches from both the domains due to the closure of the AAA1/2 cleft. The three structures in the bottom right illustrate the AAA5 bound state (**bottom**), AAA2 bound state (**middle**), and pre-power stroke state (**top**). The ILs and the AAA1/2 cleft are highlighted by the red and orange circles, respectively. (**c**) The distance between the AAA2 ILs and the LN domain is measured by Δ_*Y*1281–*T*2026_ (**yellow**), whereas the distance between the AAA5 domain and the LN is measured by Δ_*S*1315–*D*3297_ (**cyan**).

The involvement of the ILs of AAA2 in stabilizing the post-power stroke state provides a possible explanation for why ATP binding to the AAA1/2 binding site destabilizes this state. Closing of the AAA1/2 cleft brings the ILs in proximity to the AAA1 domain, making them unavailable to interact with the N-terminus regions in the LN domain. This is illustrated in Fig. S6, which shows the dynamics associated with the distance between the LN and the AAA2 ILs, and the transition of the AAA1/2 cleft. While the AAA1/2 cleft remains open, the AAA2 ILs are found to be mostly in contact with the LN domain (Fig. S6a). When the cleft closes, however, due to the binding of ATP, these contacts are destabilized (Fig. S6b). This implies that the AAA2 ILs serve as a switch for the formation of the post-power stroke state.

#### Scenario II

We expect the binding to MT to accelerate the power stroke (*LN_B_* → *LN_S_* transition). By comparing the time-dependent increase in Δ_*A*1333–*N*2341_ in the two lower panels in Fig. S7, we conclude that the LN-AAA3 detachment rate is considerably faster when dynein is bound to MT. In addition, the upper left panel in Fig. S7 also shows that in several trajectories, in which MT is bound to dynein, the LN domain reaches the AAA5 binding site (indicated by the approach of Δ_*S*1315–*D*3297_ to the value in the post-power stroke state), thus completing the power stroke. In contrast, such transitions are not observed when the motor does not interact with the MT. Therefore, we conclude that our results are in accord with the experimental observation that MT binding accelerates both ADP release, and increases the rate of the power stroke, reflected in the *LN_B_* → *LN_S_* transition.

The mechanism by which the conformational changes in the AAA1/2 cleft affect the stability of the pre-power stroke state is evident. The segment of the LN that connects the AAA3 domain to the AAA1 domain is almost perpendicular to the AAA1/2 cleft, which implies that any opening motion in the AAA1/2 cleft would pull the LN domain away from its binding sites in the AAA3 domain, thus destabilizing the pre-power stroke state. A plot of Δ_*A*1333–*N*2341_ (LN-AAA3 distance) against Δ_*N*1714–*S*2065_ (AAA1/2 cleft) in Fig. 7a and 7b shows that as the AAA1/2 cleft tends to be more open, the LN is more likely to detach from the AAA3 binding site. This effect is intensified in the MT bound state. Similarly, closing the AAA1/2 cleft would increase the stability of the pre-power stroke conformation. Thus, MT plays a key role in inducing the conformational changes in the LN. Both the opening of the AAA1/2 cleft, which occurs first, and the subsequent detachment of the LN from AAA3, occur more readily in the presence of MT.

**Figure 7:**
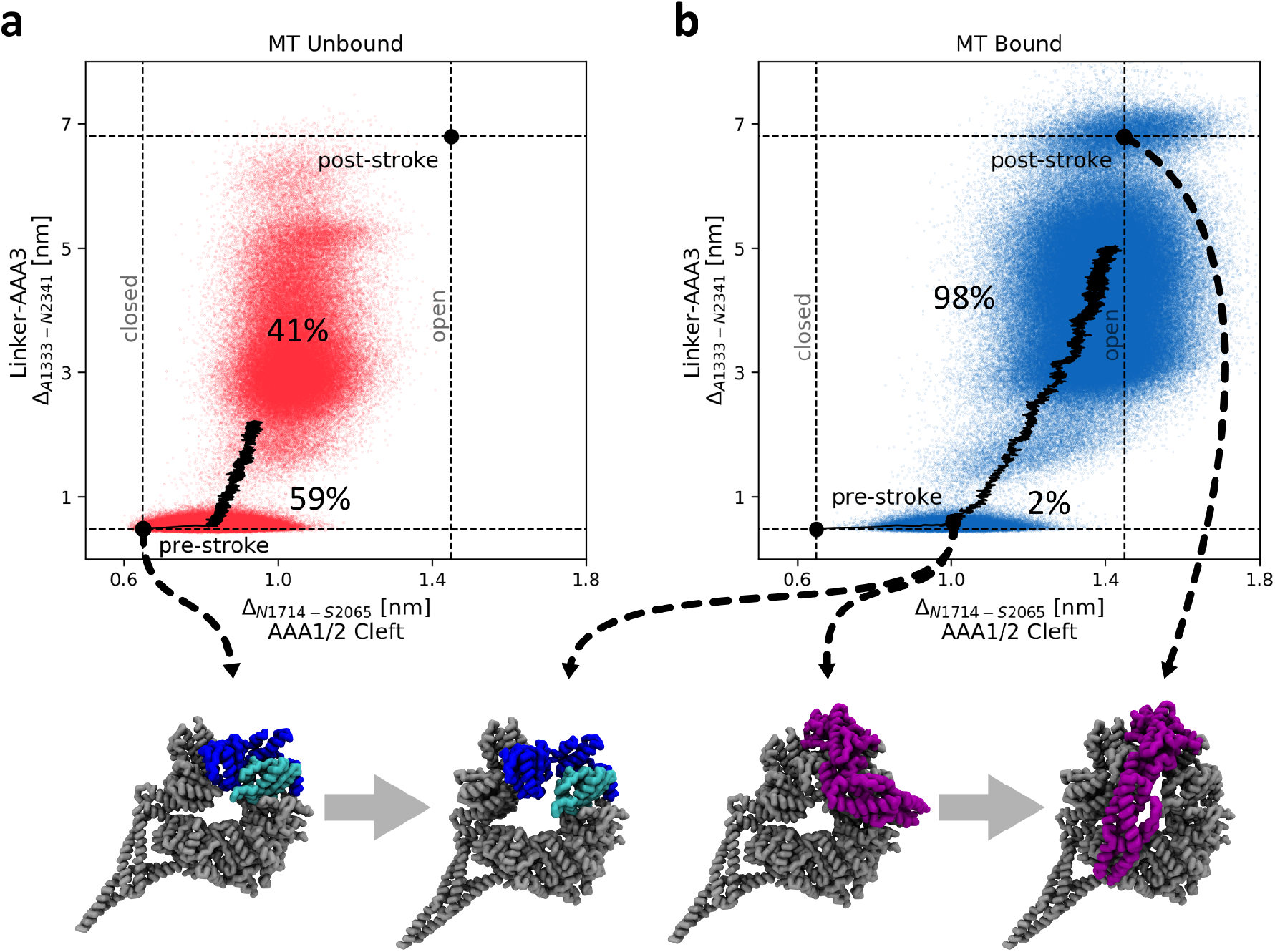
Mechanism of LN-AAA3 detachment. (**a**) Plot of Δ_*A*1333–*N*2341_ (LN-AAA3 distance) versus Δ_*N*1714–*S*2065_ (AAA1/2 cleft) generated in 100 trajectories in the absence of MT (**red**). When the AAA1/2 cleft is closed, the LN remains mostly bound to the AAA3 binding site. The percentage of trajectories in which the LN detached from the AAA3 domain, sampled during the second half of the simulation, is indicated in the plot. The mean trajectory, averaged over all trajectories, is shown by the black curve. (**b**) Variations in Δ_*A*1333–*N*2341_ (LN-AAA3 distance) against Δ_*N*1714–*S*2065_ (AAA1/2 cleft) when dynein is bound to MT (**blue**). As the AAA1/2 cleft opens, the LN detaches from the AAA3 domain and undergoes a power stroke. Dashed lines are the Δ_*A*1333–*N*2341_ and Δ_*N*1714–*S*2065_ values in the reference structures. Comparison of (**a**) and (**b**) shows the profound effect MT has on the LN-AAA3 detachment dynamics. The conformational changes in the AAA1/2 cleft (**blue**/**cyan**) and the resulting transition in the LN (**purple**) are illustrated by the four structures at the bottom of the figure.

### Inactivation of Dynein Through ATP binding to the AAA3 Domain

It is well established that the AAA1/2 domain is the principal site for ATP binding and hydrolysis in dynein.^35^ Nevertheless, several studies show that nucleotide binding and hydrolysis occur at two additional sites, located in the AAA3 and AAA4 domains respectively. Recent studies have revealed that the nucleotide state of AAA3 plays a crucial role in the regulation of dynein activity.^36, 37, 53^ In particular, while the AAA3 site contains ADP, dynein behaves normally, executing processive motion. However, when the AAA3 site is in the *apo* state or is occupied by ATP, the LN domains appears to be locked in the post-power stroke conformation regardless of the nucleotide state of the AAA1/2 site.^36^ Additional studies have shown that in this repressed state, with ATP bound in the AAA3 domain, dynein maintains a high affinity for MT as well.^36, 37^ Thus, the repressed state is incapable of stepping on MT.

In order to investigate the mechanism involved in the repression of dynein activity by the AAA3 domain, we simulated Scenario I (ATP binding to AAA1) using the crystal structure of the repressed system instead of the crystal structure of the *apo* state as a reference.^36^ We monitored the motion of the LN domain in the simulations by calculating the evolution of the distance Δ_*S*1315–*D*3297_, reporting on the LN-AAA5 interaction (structure on the right in Fig. 5c) as a function of time (Fig. 8a). In contrast to the non-repressed active motor, Fig. 8a shows that in most of the trajectories, the LN domain remained bound to the AAA5 binding site even though the AAA1/2 switched to the A state. Furthermore, the AAA1/2 cleft remained open in most of the trajectories, preventing the M to U transition in the AAA5/6/S domains (left panels in Fig. S8). This finding is consistent with the experimental results of Bhabha *et al*.^36^

**Figure 8:**
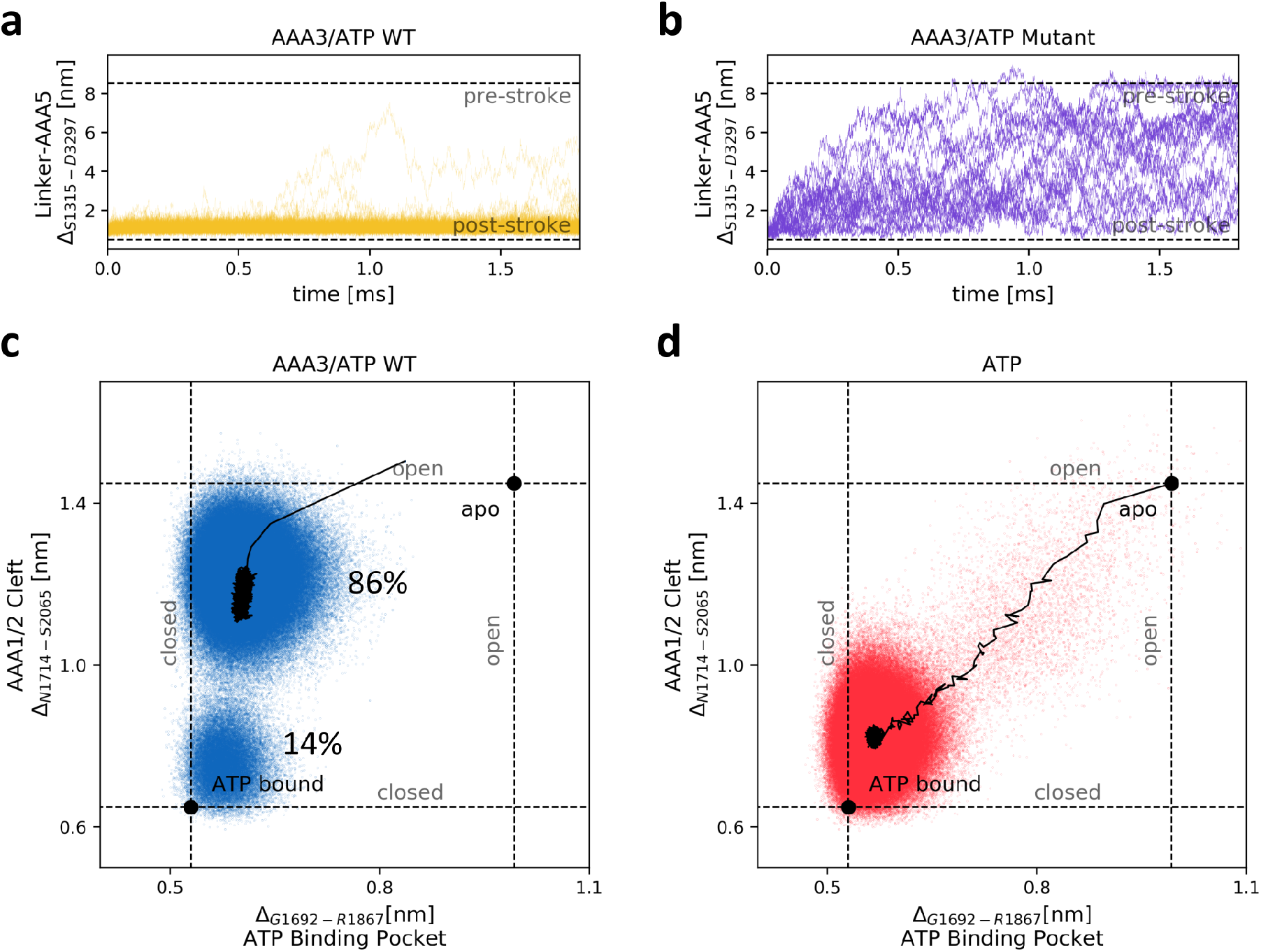
ATP bound to the AAA3 domain represses dynein activity. (**a**) The time dependent response of the LN to ATP binding in the repressed state is tracked using Δ_*S*1315–*D*3297_ (LN-AAA5 distance) in 100 trajectories. The Δ_*S*1315–*D*3297_ values in the pre- and post-power stroke are represented by the dashed lines. Even when the AAA1/2 site is occupied by ATP, the LN remains bound to the AAA5 domain in the repressed state (**yellow**). (**b**) 20 trajectories of the mutated motor in which LN-AAA2 interactions are turned off show that the LN does detach from the AAA5 domain (**purple**). This implies that the LN-AAA2 contacts are essential for the repression of dynein. (**c**) Plot of Δ_*N*1714–*S*2065_ (AAA1/2 cleft) against Δ_*G*1692–*R*1867_ (ATP binding pocket) in 100 trajectories with ATP bound to both AAA1 and AAA3 sites (**blue**). In this repressed state the AAA1/2 cleft remains open even though the nucleotide binding pocket is occupied by ATP, which would normally induce the closure of the cleft between AAA1 and AAA2 (see Fig. 3). This is a result of the AAA2 ILs contacts with the LN. The percentage of trajectories in which the cleft remained open/closed, sampled during the second half of the simulation, is indicated in the plot. The mean trajectory, averaged over all trajectories, is shown by the black curve. (**d**) Same as (**c**) except ATP is bound only to AAA1 and dynein activity is not repressed (**red**). In this state both the ATP binding pocket and the AAA1/2 cleft close due to the binding of ATP. The Δ_*N*1714–*S*2065_ and Δ_*G*1692–*R*1867_ values in the open and closed states are given by dashed lines.

We already showed that the involvement of the AAA2 domain is required in the stabilization of the post-power stroke state (Fig. S5). This is further supported by structural and mutational studies showing that the ILs play an important role in the regulation of dynein activity. ^26, 36^ In particular, the ILs come into contact with the LN domain in the repressed structure.^26, 36^ To better understand the role that the ILs play in the repression mechanism, we performed simulation of a mutated system in which the stabilizing interactions between the ILs and the LN were replaced with repulsive interactions, causing destabilization. Fig. 8b shows that in the absence of attractive LN-IL interactions the LN does detach from the AAA5 domain. Interestingly, when these interactions were turned off, the AAA1/2 cleft did close, even though the AAA3 domain was in the ATP bound state (right panels in Fig. S8). Thus, favorable interactions between the LN and AAA2 are needed to maintain dynein in the repressed state. We investigated the repression mechanism further by plotting Δ_*N*1714–*S*2065_ (AAA1/2 cleft) against Δ_*G*1692–*R*1867_ (AAA1/2 ATP binding site) in both the repressed and the nonrepressed states (Fig. 8c and 8d). Interestingly, the ATP binding site itself does close as a result of switching to the ATP bound state, even when the AAA3 domain is in the repressed state (Fig. 8c). However, the AAA1/2 cleft is open in the majority of trajectories. Because closure of the cleft is required for the formation of the pre-power stroke state the LN is unable to make a transition to the bent conformation in the repressed state. Therefore, our simulations support the hypothesis that the IL in AAA2 are crucial for the repression mechanism. In addition, the results lead to the prediction that mutations that destabilize IL-LN interactions could effect dynein motility even if it is in the repressed state.

### AAA2 ILs (Insert Loops) Involvement in Motor Gating

In order to provide structural insights into how gating could work, we performed simulations of dynein in the repressed state with ATP bound to AAA1 (scenario I), while applying a constantly increasing force at the N-terminus of the LN. We performed two sets simulations. In one set, the direction of the force was in the negative direction along the MT axis (assisting force), and in the second set, the direction of the force was along the positive direction (resisting force) (Fig. S9). As the magnitude of the force increased in both sets of simulations, the AAA1/2 cleft closed, indicating that the allosteric communication between the AAA1/2 site and the AAA5/6/S pathway was no longer repressed (Fig. S9). Interestingly, detachment of the LN from the AAA2 ILs was not required for the AAA1/2 cleft to close (Fig. 9a and 9b). The criteria for the closing of the AAA1/2 cleft was detachment of the LN from the AAA5 binding site (Fig. 9c and 9d). This suggests that the AAA1/2 cleft can close even when the LN is bound to the ILs as long as it is not simultaneously bound to the AAA5 domain. The release of LN from the AAA5 site allows it to assume conformations that can accommodate the closing of the AAA1/2, thus allowing the allosteric communication pathway to function normally. Our simulations show that both assisting and resistive force can facilitate closing of the AAA1/2 cleft even if dynein is in the non-functional repressive state.

**Figure 9:**
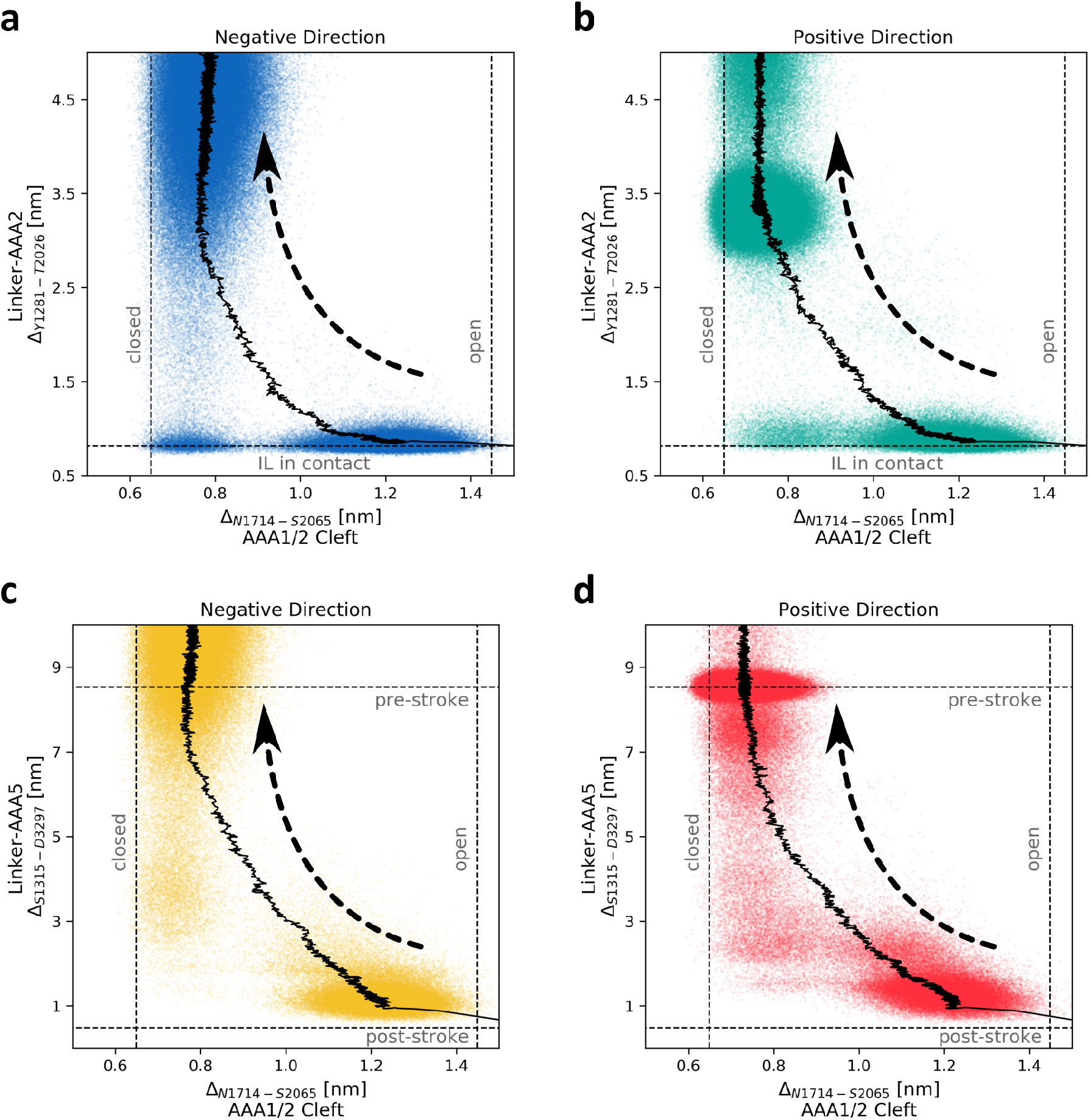
LN interactions under external load. (**a**/**b**) Plot of Δ_*Y*1281–*T*2026_ (LN-AAA2 distance displayed in Fig. 5c) against Δ_*N*1714–*S*2065_ (distance in Fig. 2b reporting conformational change in the AAA1/2 cleft) in the transitions in dynein in the repressed state (ATP bound to AAA3 domain) with increasing force applied in the negative (**blue**) and positive (**green**) directions. The mean trajectory, averaged over all trajectories, is shown by the black curve. The arrow indicates the general direction of the transition. (**c**/**d**) Same as (**a**/**b**) except these panels show Δ_*S*1315–*D*3297_ (LN-AAA5 distance) versus Δ_*N*1714–*S*2065_ (AAA1/2 cleft) with increasing force applied in the negative (**orange**) and positive (**red**) directions. While the external force causes detachment of the LN from the AAA5 domain (**c**,**d**), the LN-AAA2 contact remains intact (**a**,**b**).

We calculated the distribution of the force required to allow for the closure of the AAA1/2 cleft (Fig. S10). The histogram in Fig. S10 is an indicator of the unbinding force of the motor from the MT. In both simulation sets we obtained mean unbinding forces of the order of ≈ 40*pN*, which is about an order of magnitude larger than the unbinding forces measured in experiments.^37, 53, 54^ We attribute this to the approximations made in constructing the CG model. Interestingly, there was no significant difference between the two sets of simulations in terms of the mean value of the unbinding force. Single molecule unbinding assays indicate that the mean unbinding force tends to depend on the directionality of the applied force.^37, 53, 54^ However, this asymmetrical behavior presents itself even when the force is applied to the C-terminus of the motor, suggesting that the asymmetry is not LN dependent.

## 4 Discussion

### Reaction Cycle of Dynein

The scenarios I and II, as described above, are consistent with the common understanding of how dynein reacts to the binding of ATP and MT.^29, 38, 41^ While the reaction cycle of dynein in the repressed state is still not fully understood, a clearer picture is starting to emerge.^36, 37, 53^ Our view of the possible dynein reaction pathways, which is fully in accord with experiments, is summarized in Fig. 2 and Fig. 10. It becomes clear from our simulations that the ILs of AAA2 play a critical role in the allosteric communication in dynein, and play an even more important role in the repressed reaction cycle (Fig. 10).

**Figure 10:**
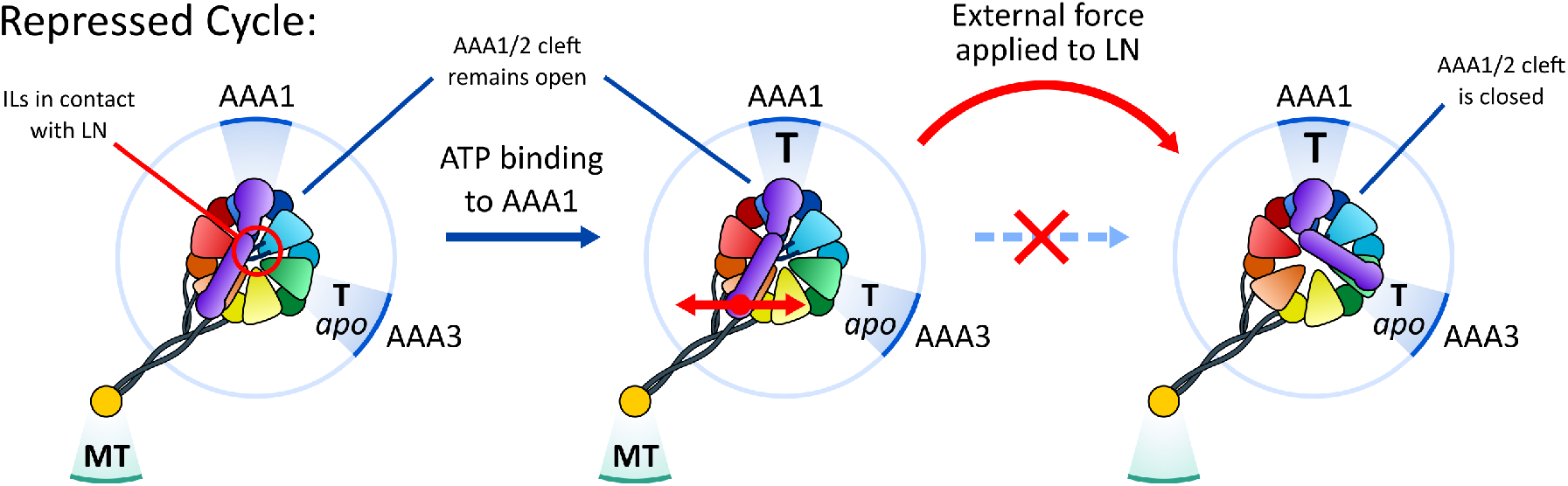
The repressed mechanochemical reaction cycle in dynein. Dynein is in the MT bound *apo* state. However, unlike in the normal cycle, the AAA3 site is either unoccupied or occupied by ATP. The ILs of AAA2 are in close contact with the LN domain (red circle in the bottom left figure). In this state the MD remains tightly bound to the MT, even after ATP binds to the AAA1 site. The AAA1/2 cleft remains partially open regardless of ATP binding. However, if an external force is applied at the N-terminus of the LN domain, the MD domain can detach from the MT and complete the priming stroke.

### ATP-Driven Formation of the Priming Stroke

Of particular interest is the way in which ATP binding to the AAA1/2 pocket translates into detachment of the LN from its binding site on the AAA5 domain, leading to the priming stroke transition. The effects of ATP binding on the AAA1/2 domains can be separated into two parts. The first is a conformational transition in the ATP binding pocket, which occurs as a result of the interactions with the bound ATP, and initiates a series of transitions across the entire AAA+ ring. The interactions with ATP are implicit in our simulations, but they are taken into account indirectly by the conformational information obtained from the crystal structure. The second part is the closure of the AAA1/2 cleft. This transition is likely due to the stabilizing interactions between the AAA1 and AAA2 domains by the conformational change in the ATP binding pocket. It is the closing motion of the AAA1/2 cleft that transmits information to the rest of the motor domain. Our simulations are consistent with the observation from experiment that changes in other domains occur only after the closing of the cleft between AAA1 and AAA2 (see Fig. 5 and S2). In the particular context of the priming stroke, closing of the AAA1/2 cleft pulls the AAA2 domain towards the AAA1 domain and the IL away from the LN. This leads to destabilization of the post-power stroke conformation as the AAA2 ILs are no longer in contact with the LN. The stabilizing interactions between the LN and the AAA5 domain are not strong enough for the two domains to stay in contact and the LN eventually undergoes a priming stroke, leading to the bent structure. Our simulations follow a hierarchy of times scales that control the conformational changes leading to the formation of the priming stroke: closing of the AAA1/2 followed both by a rapid transition in the AAA5/6/S and detachment of the ILs from the LN, leading to detachment of the LN from AAA5.

### Dynamics of the LN Conformational Transitions

A particularly interesting conclusion from our simulations is that there is a great deal of heterogeneity in the conformations of the LN domain during both the power stroke and the priming stroke (see Fig. 5, S4, and S7). Rather than taking a direct path from AAA3 to AAA5 (and vice versa), the LN assumes many orientations before binding to its final destination (Fig. S4a). Once the LN reaches either binding site (AAA3 and AAA5), however, it remains tightly bound. This is consistent with the observation from experiments that the LN assumes particular orientations in the pre- and post-power stroke states.^23, 30, 39^ Interestingly, we find that the LN spends a significant amount of time bound to the AAA2 ILs (Fig. 6), indicating that the ILs are important for the stabilization of the post-power stroke conformation. This is further supported by our simulations of the *apo* state with a mutation in the ILs. In this simulations we switched the ILs-LN interactions from attractive to repulsive, resulting in destabilization of the post-power stroke state (see Fig. S5). Our findings are supported by the observation made by Kon *et al* that when a series of mutations is introduced to the ILs, the LN assumes an intermediate conformation between the pre- and post-power stroke states, just as we observe.^26^

### Insert Loops in Dynein Motility

The fact that the AAA2 ILs are imperative for dynein proper function is appreciated.^25, 26, 36^ The precise mechanisms, however, used by the ILs to regulate motor activity have not been fully solved. Our simulations present a picture of the possible ways in which the AAA2 domain is involved in allosteric communication in the MD. We surmise that there are three important roles of the AAA2 IL. (i) Stabilizing the LN-AAA5 interactions in the *apo* state. (ii) Preventing the closure of the AAA1/2 cleft in the repressed state. (iii) Serving as an external force sensor in the repressed state (see below).

### Persistence of LN-AAA2 Interactions in the Repressed State

The LN-AAA2 interactions also play an important role in the repression of the allosteric pathway along the AAA1/2 and AAA5/6/S domains by the AAA3 unit. While it is clear that the conformational changes in the AAA3 domain are involved in the repression of dynein activity, the dynamical manner in which they do so is not clear because the changes are subtle.^36^ Our simulations reveal that AAA3 unit suppresses the detachment from MT by over-stabilizing the interactions between the LN and the ILs in AAA2 (see Fig. S8). In the crystal structure of the repressed state, the AAA3 and AAA4 domains are slightly shifted relative to one another, when compared with the other structures. This shift is sufficient to bring the AAA2 ILs closer to the LN. In addition to stabilizing the LN-AAA2 interactions, this shift may directly or indirectly destabilize the AAA1 and AAA2 interactions in the AAA1/2 cleft. As a result, the cleft is prevented from closing, even when ATP is bound at the AAA1/2 binding pocket. As shown in Fig. 6, the LN and AAA2 ILs do come into contact regardless of the nucleotide state of the AAA3 domain. However, these interactions by themselves are not enough to prevent conformational changes in the AAA1/2 domains without contribution from the AAA3 domain. This can be seen from our simulations of the ATP bound dynein in the non-repressed state (Fig. 4a and 6b), in which the AAA1/2 cleft is able to close despite the LN interactions with the ILs.

### Mechanical Forces Rescue the Motility of Dynein in the Repressed State

Studies show that even when the AAA3 domain is occupied by ATP, if a strong enough external mechanical force is applied at the N-terminus of the LN domain, dynein resumes normal activity,^37^ thus restoring processivity. The results of our simulations, which provide a molecular picture of communication between AAA1 and ATP-bound AAA3 needed for stepping, are consistent with experiments. Interestingly, we find that pulling on the LN does not immediately destabilize the interactions of the LN with the AAA2 domain (Fig. 9a-b). Instead, breaking the contacts between the LN and the AAA5 domain appear to be sufficient to reverse the mobility suppressing effects of the AAA3 when dynein is the repressed state (Fig. 9c-d). This implies that the combination of LN interactions with both the AAA2 and AAA5 domains as well as the stabilization of these interactions by the ATP bound AAA3 domain are required for repression of dynein activity. It is sufficient to disrupt interactions associated with one of these three elements to allow dynein to undergo the priming stroke and detach from MT. As we mentioned earlier, there have been previous studies of dynein in which the AAA2 ILs were mutated.^26, 36^ However, these studies focused on the effects of mutations on the ATPase activity and LN conformation but did not measure their effect on the affinity of dynein for MT. We propose that force dependent unbinding experiments on dynein, such as those reported recently^37^ with mutations of the AAA2 IL, be performed to test our predictions regarding the role of the AAA2 IL in the repression mechanism.

### Effect of Mechanical Force and Implications for Gating

In the context of gating, it is worth pointing out that in our simulations, pulling in both the negative and positive directions along the MT axis had approximately the same effect. This implies that the AAA3 repression system does not work as a gating mechanism in the classical sense of creating an asymmetry between the leading and trailing MDs. Instead, it prevents dynein detachment from the MT in the absence of external strain, indicating that the repressed state, in addition to potential regulation mechanism may play an important in the context of processivity of dynein.^36^

Since we know that dynein does respond asymmetrically to an external load, it might seem that our findings may not be correct.^54^ This is, however, not necessarily the case as there are other possible reasons for the asymmetric response. One reason could be that when dynein binds to MT, it does so at an angle.^55, 56^ This, by itself, could be sufficient to elicit an asymmetric response to force, especially in the context of single molecule unbinding experiments.^37, 53, 54^ It has been pointed out that when the MDs in the dynein dimer are far apart, there is a higher probability that the trailing MD is likely to step.^57, 58^ This indicates that there is some gating mechanism in dynein that is sensitive to internal strain. Our results do not account for this effect, because we have focused on transitions that occur in a single motor. A reasonable explanation, however, can be given by the possibility that a backward load, acting on the leading MD, would either prevent or slow down the power stroke associated with the leading motor, which is necessary for ADP release and completion of the dynein mechanochemical cycle. This is a plausible gating mechanism that does not require an asymmetric response to force, and would be consistent with our findings. We hasten to add that until simulations of the dynein dimer are performed, preferably in complex with MT, we cannot clarify the origin of the observed asymmetric response to tension. However, the structural reasons for restoration of dynein motility upon application of external force, predicted here, is likely to be robust.

## 5 Conclusions

By taking advantage of the available structures of dynein in various nucleotide binding states, we constructed a new coarse-grained model to investigate the dynamics of allosteric transitions in cytoplasmic dynein, and performed Brownian dynamics simulations. The model vividly illustrates the hierarchy of structural changes that occur when ATP binds to AAA1 and propagate through the entire MD, thus producing the pre-power stroke state with the LN domain in the bent conformation. Although the observed series of structural transitions was already anticipated by comparing the end states of dynein from several structures in different nucleotide states, our simulations provide some estimates of the relative rates and molecular changes that occur as a result of binding of ATP to AAA1.

Because our simulations monitor the dynamics of the various allosteric transitions, we are able to assess the order of structural transitions. For example, the scatter plot in Fig. 6 shows that only after the LN forms contacts with the ILs of AAA2 does the LN detach from AAA5. Similar time-dependent changes that must occur during other allosteric transitions naturally emerge from our simulations. Our work supports the observation that even the repressed state could be rescued by application of an external force. The main prediction is that the insert loops in AAA2 play a crucial role in the repression mechanism as well as in the ability of dynein to respond to external force. The prediction that alteration of interactions between ILs and AAA2 could impact motility in the repressed state even when the LN is not subject to a mechanical force is amenable to experimental tests. Finally, the present work showcases the efficacy of coarse-grained models in providing the dynamics of allosteric transitions in complex molecular machines such as dynein.

## 6 Experimental Procedures

### Self Organized Polymer Model

In order to monitor the allosteric transitions in the dynein MD we performed Brownian dynamics simulations, using the SOP model, which has proven to be remarkably successful in modeling the behavior of motor proteins.^43, 44, 46, 59^ In the SOP representation each amino acid is represented by a single bead, centered at its *C_α_* carbon. The energy function that governs the interactions between the different beads is described in detail in the SI.

### State-Dependent Triggers of the Allosteric Transitions

A fundamental aspect of our approach is that the allosteric transitions in dynein occur only in response to external stimuli. More specifically, the priming stroke and detachment from the MT occur in response to the binding of ATP to the AAA1 domain. The power stroke and ADP release are consequences of rebinding to the MT. This behavior is introduced in our model by using the conformations of the AAA1/2 cleft (ATP binding) and the AAA5/6/S domains (MT binding) as the systems inputs. The conformational changes of the LN domain, which occur in response to the conformational transitions in these domains, are the outputs that are automatically produced in our simulations.

In order to simulate the effects of the binding of ATP to the AAA1 domain, we stabilized the ATP bound, A state of the AAA1/2 domain (upper row in Fig. 2). Only after the AAA1/2 cleft is closed, the conformational transition of the AAA5/6/S domains from the M state to the U state was initiated. Similarly, in order to simulate the binding of dynein to the MT, in the ADP bound state, we stabilized the M state in the AAA5/6/S domains (lower row in Fig. 2). The AAA1/2 was allowed to transition to the open E state after the transition of the AAA5/6/S domains is complete.

In order for the LN to respond spontaneously to the conformational changes of the AAA1/2 domain, we used a double-well SOP potential, as described by the third term of Eq. 2 in the SI. This allows the LN to transition freely between the bent *LN_B_* state and the straight *LN_S_* state. Therefore, the particular conformations adopted by the LN in our simulations are entirely due to a response to the conformational changes of the AAA1/2 and AAA5/6/S domains.

## Acknowledgements

We are grateful to the National Science Foundation for supporting the work through grant CHE 16-36424. Additional support from the Collie-Welch Chair (F-0019) is greatly appreciated.

